# Tissue-specific tagging of endogenous loci in *Drosophila melanogaster*

**DOI:** 10.1101/025635

**Authors:** Kate Koles, Anna R. Yeh, Avital A. Rodal

## Abstract

Fluorescent protein tags have revolutionized cell and developmental biology, and in combination with binary expression systems they enable diverse tissue-specific studies of protein function. However these binary expression systems often do not recapitulate endogenous protein expression levels, localization, binding partners, and developmental windows of gene expression. To address these limitations, we have developed a method called T-STEP (<u>T</u>issue-<u>S</u>pecific <u>T</u>agging of <u>E</u>ndogenous <u>P</u>roteins) that allows endogenous loci to be tagged in a tissue specific manner. T-STEP uses a combination of efficient gene targeting and tissue-specific recombinase-mediated tag swapping to temporally and spatially label endogenous proteins. We have employed this method to GFP tag OCRL (a phosphoinositide-5-phosphatase in the endocytic pathway) and Vps35 (a Parkinson's disease-implicated component of the endosomal retromer complex) in diverse *Drosophila* tissues including neurons, glia, muscles, and hemocytes. Selective tagging of endogenous proteins allows for the first time cell type-specific live imaging and proteomics in complex tissues.

## Introduction

Cellular and developmental biology has been transformed by the application of fluorescent tags, enabling the localization and live imaging of specific proteins and biochemical isolation of their binding partners among a large number of diverse applications. In *Drosophila melanogaster*, the introduction of the first binary UAS-Gal4 system in 1993 (Brand and Perrimon 1993) allowed for the tissue specific expression and analysis of proteins of interest, including fluorescently tagged proteins. Even though the UAS-Gal4 (and other binary expressions systems) are indispensable in any *Drosophila* laboratory, in some experimental contexts they do not recapitulate endogenous protein levels and regulatory elements. In these scenarios, toxicity, non-physiologically relevant protein localization or activity can often arise due to artificially high protein expression levels, or by ectopic expression in tissues where the gene of interest is not naturally expressed.

Given our interest in identifying the native localization pattern and binding partners of endocytic proteins in a tissue-specific manner, we sought to eliminate these common shortcomings while preserving the tissue-specificity of the binary expression systems. We designed our method to be economical and easily adopted by any laboratory. By combining the highly efficient lethality selection based gene targeting approach (Chen et al. 2015) with a recently introduced recombinase R (which we refer to as Rippase) from the yeast *Zygosaccharomyces rouxii* (Nern et al. 2011) here we demonstrate the efficiency and effectiveness of the T-STEP method to tissue-specifically label any protein, allowing for cell type-specific imaging and biochemical analysis at endogenous levels.

## Materials and Methods

### DNA constructs

Standard molecular biology techniques and Gibson cloning were used to generate all plasmids and intermediates. The pT-STEP donor plasmids for C-terminally tagged GFP-swappable TagRFPT fusions incorporate the recently published lethality selection (from vector pTL2 (Chen et al. 2015)) with the following modifications: The TagRFPT coding region was amplified from TagRFPT-EEA1, (Addgene plasmid #42635, from Silvia Corvera) with primers incorporating the Rippase recognition sequence RRS (in -1 frame relative to the directionality of RRS so that no stop codons are present: <u>TTGAT GAAAGAAT ACG TT ATT CTTT CATCAA</u>) in frame at the 5' of TagRFPT, leading to a short linker peptide when translated (LMKEYVjLSS-S-TagRFPT). The 3'-UTR from the *Autographa californica* nucleopolyhedrovirus (AcNPV) p 10 gene was amplified from pJFRC81 -10XUAS-IVS-Syn21-GFP-p 10, (Addgene plasmid # 36432, from Gerald Rubin), chosen for its efficiency in female germline cells (Pfeiffer, Truman, and Rubin 2012). The LoxP-DsRed-LoxP cassette was obtained from pDsRed-attP (Addgene plasmid # 51019, from Melissa Harrison, Kate O'Connor-Giles and Jill Wildonger). The PreScission Protease (PSP) recognition sequence (TTGGAGGTCCTGTTCCAGGGCCCC/LEVLFQ^GP) followed by a short linker (GSGSGS) and GFP were synthesized as a gene block (IDT DNA, Iowa). The second RRS recognition sequence (-1 frame) was included in the 5' primer used to amplify the PSP-GFP gene block.

The BspQI sites within pTL2 were changed to BsmBI (TagRFPT contains a BspQI site). Components of the swappable-to-GFP cassette were first assembled in a PCR4-TOPO vector yielding the intermediate pDsRed-TSTEP vector (which could serve as a starting vector for laboratories preferring injection based gene targeting, see also Supplemental Figure 4). Using this intermediate as a template, the RRS-TagRFPT-p10-LoxP region was PCR amplified and inserted between the NsiI and SpeI of pTL2, thereby removing the original I-CreI and attPX sites. The resulting plasmid was then digested with PacI and BamHI and, using Gibson cloning, the RRS-PSP-GFP was inserted yielding the final pT-STEP vector, which has been deposited at Addgene. The pT-STEP-SNAPf vector is identical to pT-STEP except that a SNAPf tag (New England Biolabs) instead of the GFP tag is introduced after the Rippase reaction (see also Supplemental Figure 4). All vector details are available on request. Oligoes corresponding to CRISPR Cas9 target sites (Vps35 [t] agcccagcgcacccactt and OCRL [c]cgcagctgtgccgccgaat) and containing 4 extra base pairs for BsmBI compatibility (see Supplemental Figure 5) were annealed and ligated the BsmBI site of pT-STEP (bases in [brackets] were changed to the obligatory G for the dU6.3 promoter). For introducing the Parkinson's disease specific human Vps35D^620^N mutation (corresponding to D628N in *Drosophila melanogaster* Vps35), the 5' arm of wild type Vps35 was subcloned into pJet1.2 vector and Gibson cloning was used to introduce the specific mutation. Three targeting vectors were made (OCRL, Vps35 and Vps35D^628^N). The 5' homology arms of wild type or D628N mutant Vps35 (2R:22I85904..22I89272) or OCRL (X:1924260.. 1927163) were inserted at the StuI site of pT-STEP, and their respective 3' arms (2R:22189273..221 92812) and (X: 1927164..1930079) in the PmeI site (numbers reflect the DGRC r6.05 database). The Vps35 3' homology arm was modified to abolish the PAM region of the chosen target sequence, while the OCRL targeting construct did not carry resistance to Cas9.

#### Gene targeting with lethality selection (Golic+)

The donor plasmid for OCRL was injected to attP40 docking site and for Vps35 to the VK00027 docking site by Rainbowgene Inc, CA. All Golic+ strains were from Hui-Min Chen and Tzumin Lee (Janelia Farms) (Chen et al. 2015), and the bam898-Cas9-2A-FLP-2A-ISceI stocks were generously shared before publication (See also Supplemental Figure 5 for detailed workflow). Transformants were screened for by crossing to the Pin/CyO; GMR3-LexA in attP2 and GMR3-LexA in attP40; TM3, Sb/TM6B, Tb, respectively, using the reduced eye size phenotype. The Vps35 donor lines (Vps35-790.25.F1 in VK00027and Vps35^D628^N-795.46.M4 in VK00027) were crossed to bam898-Cas9-2A-FLP-2A-I-SceI in su (Hw)attP8/FM6c; Pin/CyO; LexAop2-5xriTS-RacI^V12^ in VK00027/ TM3, Sb strain while the OCRL donor line (OCRL-793.R39.1 in attP40) to the LexAop2-5xriTS-RacI^V12^ in attP40/CyO; bam898-Cas9-2A-FLP-2A-I-SceI in attP2/TM3, Sb. Lethality selection was then performed using the Pin/CyO; nSyb-LexA in VK00027 for Vps35 and FM7a; nSyb-LexA in attP16 for OCRL. Candidates were mapped using the Pin/CyO; GMR>riTS-RacI^V12^ line.

#### Antibodies

FasII ID4 (I:10), LamDm0 (1:500) and LamC (1:30) monoclonal antibodies were from Developmental Studies Hybridoma Bank (University of Iowa). Anti-TagRFP antibody (1:1500 for Western blotting) was from Evrogen (AB234). Anti-HRP antibodies and secondary antibodies for imaging were conjugated to Alexa 488 or Alexa 647 (Jackson Immunoresearch). Anti-rabbit DyLight 680 antibody (I:5000) was from Rockland, PA.

#### Hemocyte preparation and imaging

Hemolymph from third instar larvae was bled onto coverslips into a drop of phosphate buffered saline (PBS) containing 20 μM phenyl-thiourea (PTU) and allowed to adhere for 5 min at room temperature. Hemocytes were either imaged live by mounting the coverslip on a glass slide with narrow spacer, or fixed in 4% paraformaldehyde on the coverslips in PBS for 10 min and then stained with the indicated antibodies.

#### Fly strains

Flies carrying the UAS-Rippase followed by a short PEST sequence (aa 422—461 of the mouse ornithine decarboxylase gene) to decrease its half-life and potential toxicity (pJFRC165-20XUAS-IVS-R::PEST in attP2) were from Gerald Rubin (Janelia Farms); Alrm-Gal4 was from Marc Freeman (UMASS Medical School), C380-Gal4 and C57-Gal4 were from Vivian Budnik (UMASS Medical School), UAS-GFP-Rab5 from Marcos Gonzalez-Gaitan (University of Geneva, Switzerland) and GFP-Rab5 knock-in flies (Fabrowski et al. 2013) from Stefano De Renzis (European Molecular Biology Laboratory (EMBL) Heidelberg, Germany). YFP-HA-RabI I knockin flies (Dunst et al. 2015) were from Marko Brankatschk (Max Planck Institute of Molecular Cell Biology and Genetics, Germany), Hem-ese-Gal4 (#8699), ddc-Gal4 (#7009), Repo-Gal4 (#7415) and w; Df (exel)6078/CyO (#7558) were from the Bloomington Drosophila Stock Center.

#### Western blotting

Hemolymph from ten third instar larvae was collected into 10 μl of 20 μM PTU in PBS as illustrated in https://www.youtube.com/watch?v=im78OIBKlPA. An equal volume of 2x sample buffer (BioRad) was added and samples were heated for 5 min at 95 oC. 15 μl (3.8 larvae) were loaded onto 7.5% polyacrylamide gels (Bio-Rad). Proteins were transferred to nitrocellulose membrane and blocked in blocking buffer for fluorescent western blotting (Rockland, PA). The membrane was probed with rabbit anti-TagRFP antibodies followed by anti-rabbit DyLight 680, and visualized on Licor Odyssey Scanner.

#### Larval dissections and imaging

Third instar larvae were dissected in HL3.1 (Feng, Ueda, and Wu 2004) and fixed in 4% paraformaldehyde in HL3.I for I0 min at room temperature, then rinsed and stained with appropriate antibodies in PBS containing 0.2% (v/v) TritonX-100. Larvae were mounted in Vectashield (Vectoi Labs, Burlingame, CA). Spinning-disk confocal Z-stacks (0.3 μm or 1 μm were collected at room temperature on an Andor spinning-disk confocal system consisting of a Nikon Ni-I upright microscope equipped with 40 X (numerical aperture [NA] 1.3) 60 X (NA 1.4) and 100 X (NA 1.45 oil immersion objectives, or a 60 X (NA 1.0) water immersion objective a Yokogawa CSU-W1 spinning-disk head, and an Andor iXon 897U electron-multiplying charge-coupled device camera (Andor, Belfast, Northern Ireland). Images were collectec using Nikon Elements AR software and processed using Volocity software (Improvision).

### Results and Discussion

#### Rationale for T-STEP

Binary expression systems in *Drosophila*, such as the UAS-Gal4, LexAop-LexA and QUAS-QF, offer tissue-selective visualization and manipulation of genes of interest. However, these methods do not faithfully recapitulate endogenous protein expression levels and/or localization. An example of such an effect and the dramatic improvement that can be achieved by genomic tagging is shown in Figure 1 at the *Drosophila* third instar larval neuromuscular junction (NMJ). In this example, the endogenously GFP-tagged Rab5 protein (Fabrowski et al. 2013), a marker for early endosomes, exhibits very different localization from an UAS-GFP-Rab5 transgene expressed with the neuronal C380-Gal4 driver. While the endogenous GFP-Rab5 localizes to small, fairly uniform puncta, in both the motor neuron and in surrounding muscle tissue, neuronally overexpressed GFP-Rab5 is concentrated in enlarged compartments. Thus, overexpression of Rab5 dramatically changes its localization. While endogenous gene tagging resolves these overexpression issues, it does not enable biochemical approaches to selectively isolate the interacting partners of Rab5 at endogenous protein levels from specific tissue types (such as muscle, glia or neurons) since all tissues express the same GFP-tagged Rab5. Furthermore, live imaging at endogenous levels in complex, intertwining tissues (such as glia and neurons) is also challenging due to spatially overlapping signals. We addressed these limitations by designing a gene targeting cassette, T-STEP (<u>T</u>issue-<u>S</u>pecific <u>T</u>ag-ging of <u>E</u>ndogenous <u>P</u>roteins), comprised of two key components, a) tandem Rippase specific recognition sequences (RRS) in frame with the targeted protein, which allows tissue-specific tag switching and b) a lethality selection cassette for very high efficiency gene targeting (Chen et al. 2015) (Figure 2 and Supplemental Figure 1 to Figure 2). Recombinase R, or Rippase, was identified in yeast *Zygo-saccharomyces rouxii*, and it is one of four novel site-specific recombinases recently adopted in flies (Nern et al. 2011). Like other recombinases, Rippase mediates extremely efficient (>96%) DNA exchange between two Rippase specific recognition sequences (RRS), and is fully compatible with other existing genetic tools such as FLP/FRT. Most relevant for the T-STEP method, the recognition target sequence of Rippase (RRS, blue arrows in Figure 2 and Supplemental Figure 1 to Figure 2) can be translated without stop codons, and when in frame with the tagged protein, it serves as a short peptide linker between the C-terminus of the targeted protein and the TagRFPT or GFP tag (Fig 2 and Supplemental Figure 1 to Figure 2B and C). Another crucial component of our approach is the extremely efficient lethality selection cassette adapted from (Chen et al. 2015), without which T-STEP would not be easily accessible for many fly labs. Compared to all existing gene targeting methods (Gratz et al. 2015; Gratz et al. 2014; Zhou et al. 2012) that require molecular or visual screening of often very large numbers of gene targeting candidates, the novel and innovative design of lethality selection kills flies bearing non-or mis-targeted events, leaving only correctly targeted flies viable, thereby eliminating the need for labor intensive screening (for detailed information on the design features of lethality selection see (Chen et al. 2015)). Furthermore, since the location of dsDNA break is highly restricted to the gene region targeted by T-STEP (Supplemental Figure 1 to Figure 2B), the number of available gRNA target sequences might be limited, which may necessitate the use of gRNA sequences with low efficiencies. Lethality selection easily compensates for potentially low-efficiency gRNAs by simply scaling up the number of crosses without any extra effort at injection or screening. Thus lethality selection allows any laboratory without access to large-scale embryo injection facilities to target any gene with the T-STEP cassette in a virtually fail-proof manner, with unprecedented ease and speed (see (Chen et al. 2015)). We have also generated a 3xP3-dsRed marked version of the T-STEP vector for labs preferring injection based gene targeting with visual screening for targeted events (Gratz et al. 20I4) and Supplemental Figure 4.

**Figure 1. Overexpression of the endosomal marker GFP-Rab5 changes its localization and distribution pattern.**
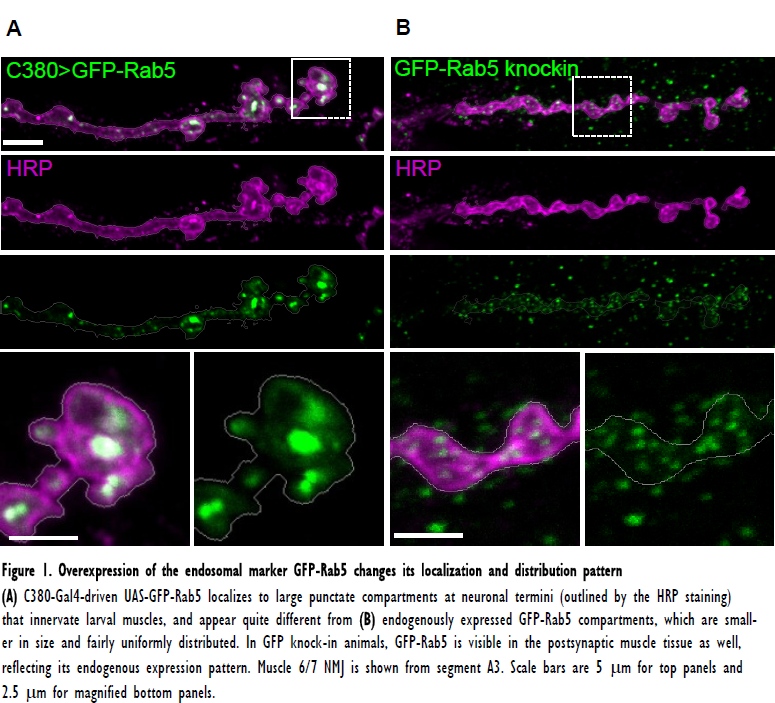
**(A)** C380-Gal4-driven UAS-GFP-Rab5 localizes to large punctate compartments at neuronal termini (outlined by the HRP staining) that innervate larval muscles, and appear quite different from **(B)** endogenously expressed GFP-Rab5 compartments, which are small-er in size and fairly uniformly distributed. In GFP knock-in animals, GFP-Rab5 is visible in the postsynaptic muscle tissue as well, reflecting its endogenous expression pattern. Muscle 6/7 NMJ is shown from segment A3. Scale bars are 5 μm for top panels and 2.5 μm for magnified bottom panels.

**Figure 2. Conceptual design of the Tissue-Specific Tagging of Endogenous Proteins (T-STEP) method.**
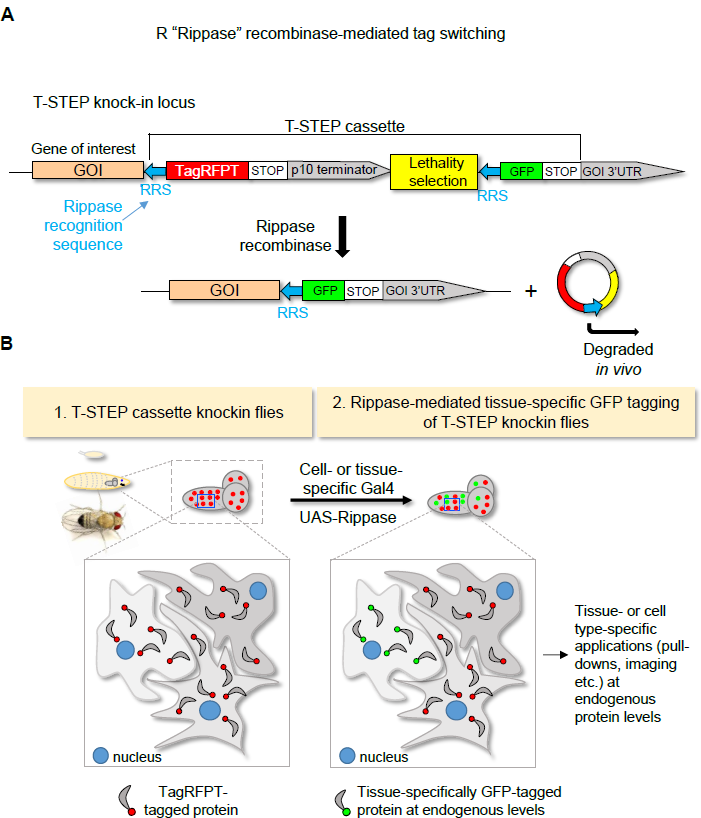
**(A)** Schematic outline of the genomic locus after T-STEP cassette knockin using gene targeting. Translation of the targeted protein yields a TagRFPT-tagged protein with the translated Rippase recognition sequence, RRS, serving as a short peptide linker. Upon tissue-specific expression of Rippase the DNA sequence between the tandem RRS se-quences is excised and degraded, rendering the protein of interest GFP-tagged and under fully endogenous 5' and 3' regulatory elements. **(B)** Before Rippase expression, the targeted protein is TagRFPT-tagged in all cells where it is naturally expressed. Tissue-or cell-type specific protein labeling is achieved by Rippase expression in the desired cell-or tissue type using the UAS-Gal4 system.

We tested our approach by T-STEP tagging wild-type Vps35 (2^nd^ chromosome) (see Supplemental Figure I to Figure 2B for details of the targeting steps) and OCRL (X chromosome). For Vps35, we made a second targeting construct that also carried the conserved Parkinson's disease-linked D628N linked mutation(Zimprich et al. 2011) in the 5' homology arm. The donor vectors were inserted into the appropriate attB docking site via standard transgenesis. Following a simple crossing scheme with published stocks ((Chen et al.2015) and Supplemental Figure 5) we obtained targeted events for all three constructs with very high efficiency (see SupplementalTable I), which were further confirmed by Westernblotting and PCR (Supplemental Figure 2).

#### In vivo characterization of T-STEP knockins

Imaging fixed third instar larval tissues demonstrated the subcellular localization of endogenous OCRL-TagRFPT and Vps35-TagRFPT protein, respectively (Supplemental Figure 3) as well as their subcellular dynamics upon live imaging (data not shown). In hemocytes, OCRL-TagRFPT localized to small, fairly uniformly distributed structures throughout the cytoplasm, likely of endocytic origin, as well as in the nucleus (Supplemental Figure 3A). Vps35-TagRFPT was expressed at higher levels than OCRL-TagRFPT, and was the focus of our remaining experiments. Vps35-TagRFPT was readily visible in in most tissues, including the nervous system, epithelia, muscles, and hemocytes, where Vps35 has previously been shown to function (Korolchuk et al. 2007, Dong et al. 2013). Live imaging of Vps35-TagRFPT in hemocytes revealed its dynamic association relative to Rab5-or RabI I-positive endosomes (data not shown). In fixed larval muscle cells Vps35-TagRFPT was found in small, distributed puncta and in larger perinuclear structures (Supplemental Figure 3 B and C). Thus, the T-STEP cassette efficiently reports the localization of targeted endogenous proteins.

#### Tissue-specific Rippase-mediated GFP tagging of T-STEP knockins

To test whether tissue-specific expression of the Rippase could lead to the conversion of Vps35-TagRFPT to Vps35-GFP, we employed a range of tissue-specific Gal4 drivers. In all tissues tested we observed the appearance of Vps35-GFP (Figures 3 and 4), in accord with the very high efficiency of Rippase mediated events reported previously (Nern et al. 2011). In a population enriched for glutamatergic motor neurons (C380-Gal4), Vps35-GFP was detected in neuronal cell bodies as well as the neuropil (Figure 3A). When we expressed Rippase using ddc-Gal4 (which expresses Gal4 in a subset of dopaminergic and serotonergic neurons (Li et al. 2000)), the Vps35-GFP signal revealed in unprecedented detail the subcellular localization of Vps35 in a tissue type implicated in Parkinson's disease (Figure 3B).

**Figure 3. Tissue-specific tagging with T-STEP.**
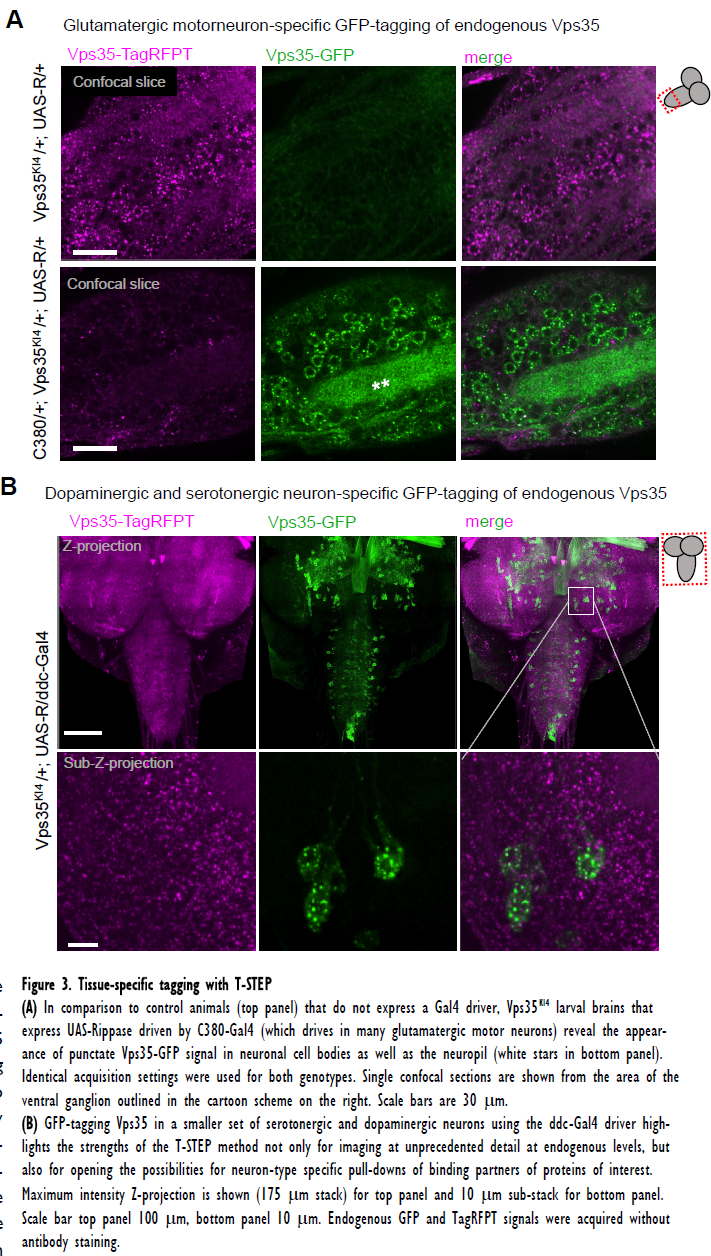
**(A)** In comparison to control animals (top panel) that do not express a Gal4 driver, Vps35^KI4^ larval brains that express UAS-Rippase driven by C380-Gal4 (which drives in many glutamatergic motor neurons) reveal the appearance of punctate Vps35-GFP signal in neuronal cell bodies as well as the neuropil (white stars in bottom panel). Identical acquisition settings were used for both genotypes. Single confocal sections are shown from the area of the ventral ganglion outlined in the cartoon scheme on the right. Scale bars are 30 μm. **(B)** GFP-tagging Vps35 in a smaller set of serotonergic and dopaminergic neurons using the ddc-Gal4 driver highlights the strengths of the T-STEP method not only for imaging at unprecedented detail at endogenous levels, but also for opening the possibilities for neuron-type specific pull-downs of binding partners of proteins of interest. Maximum intensity Z-projection is shown (175 μm stack) for top panel and 10 μm sub-stack for bottom panel. Scale bar top panel 100 μm, bottom panel 10 μm. Endogenous GFP and TagRFPT signals were acquired without antibody staining.

**Figure 4. Vps35 tagging in astrocytes, pan-glial populations, larval muscles and larval hemocytes.**
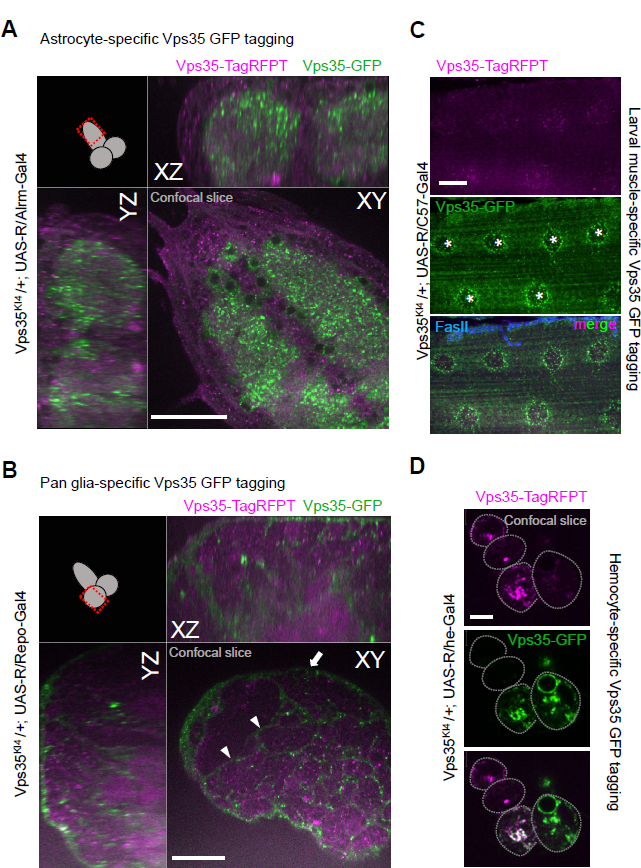
**(A)** YZ and XZ single confocal planes demonstrating the subcellular distribution of Vps35-GFP throughout the larval astrocyte population, including their processes infiltrating the neuropil. Scale bar 40 μm. **(B)** Repo-Gal4-mediated pan-glial tag conversion reveals surface glial (arrow) and cortex glial (arrowheads) expression of Vps35 in a single confocal plane through the lobe of the larval CNS. Scale bar 40 μm. **(C)** Maximum intensity projection of a confocal Z-stack through a larval muscle cell M6 (which are multi-nucleate, marked by stars) that expresses Rippase with the muscle driver C57-Gal4. Vps35-GFP is present most prominently around the muscle nuclei though it can be seen in small punctate structures throughout the muscle cell. Synaptic termini were stained with FasII. Scale bar 25 μm. **(D)** Single confocal slice through a cluster of four third instar larval hemocytes from Vps35^KI4^ knock-in animals that express Rippase with the hemese-Gal4 driver in the majority of hemocytes (Zettervall et al. 2004). The top two cells express Vps35-TagRFPT almost exclusively, with only very low level of Vps35-GFP signal detectable, indicating that Vps35-GFP translation has not progressed long enough for the signal to become readily visible. The other hemocytes accumulated varying amounts of Vps35-GFP depending on their cellular birth date and the relative timing of R-mediated conversion. Scale bar 5 μm.

When tagged in astrocytes, Vps35-GFP localized to astrocyte cell bodies as well as to processes infiltrating the neuropil (Figure 4A). Pan-glial tagging using Repo-Gal4 revealed that Vps35 is expressed in a number of diverse glia types (Figure 4B). In larval muscles, Vps35 was most prominent around the muscle nuclei (Figure 4C). In hemocytes, Vps35-GFP was readily observed in the same pattern as Vps35-TagRFPT (Figure 4D). In this tissue type (and to a lesser extent in the larval muscles) we noted some variability in the ratio of Vps35-TagRFPT to Vps35-GFP (Fig 4D) likely reflecting a combination of factors ranging from Vps35 protein half-life, strength of the Gal4 driver, tissue or cell-type specific protein levels, and the timing of the Rippase-mediated event relative to cell division. These variables of the T-STEP sys-tem could potentially be exploited to assess the half-life of proteins pre-and post-Rippase-mediated conversion in specific tissue-types or during specific developmental windows.

One potential caveat of any protein tagging system is that the tag could interfere with protein function, localization or degradation. The Vps35 and OCRL homozygous T-STEP knockin flies are fertile and viable (compared to null mutants, which are larval lethal ((Korolchuk et al. 2007) and our unpublished results) and the Vps35KI allele complements a small chromosomal deficiency lacking Vps35. The TagRFPT-tagged version of the knockins carries the p10 terminator from *Autographa californica* nucleopolyhedrovirus, while the GFP-tagged rip-out allele is under fully endog-enous regulatory elements. Thus, it is possible that their regulation might be different. Howev-er, the identical localization pattern of Vps35 and OCRL (data not shown) before and after tag conversion argues that in the case of these two proteins, the p10 3'UTR does not negatively interfere with expression patterns or localization. Although p10 was initially chosen to minimize the presence of repetitive regions in the donor vector, it should also be possible to use endogenous 3' regulatory elements for both TagRFPT and GFP tagged versions of the targeted proteins.

One of the inherent drawbacks of our approach is that it may be of limited use for proteins with very low expression levels. We have prepared T-STEP cassettes with alternative tags, such as SNAPf, which may offer further flexibility and sensitivity for certain applications (Kohl et al. 2014). In addition, T-STEP could be used to simultaneously label both an mRNA and its cognate protein in a tissue-specific manner, by incorporating RNA-tagging recognition sequences in the 3'UTR of the targeting cassette. This would allow the method to be extended for the tissue-specific identification of protein and/or mRNA binding partners at endogenous levels.

Furthermore, T-STEP offers unique opportunities to facilitate the mechanistic understanding of diverse tissue-specific diseases. For example, in many neurological diseases select neuronal populations are predominantly affected (e.g. motor-neurons in Amyotrophic Lateral Sclerosis, or dopaminergic subpopulations in Parkinson's disease), even though every cell of the organism carries the causative mutation. By using T-STEP and taking advantage of existing and rapidly expanding (Diao et al. 2015) tissue-specific drivers, one can selectively visualize, analyze or isolate protein or RNA from the affected tissues of wild type or mutant animals at native protein and RNA levels, a possibility that has not been feasible until now. In summary, the T-STEP approach affords a simple and robust method to tissue-specifically label proteins at their C-termini at endogenous levels, and with comparable cloning effort that is required for routine binary expression constructs (UAS/LexAop/QUAS).

## Acknowledgements

Stocks and reagents were generously provided by Drs. Hui-Min Chen and Tzumin Lee, Gerald Rubin, the Bloomington *Drosophila* Stock Center (NIH P40OD0I8537), and the Developmental Studies Hybridoma Bank (University of Iowa). We thank Drs. Hui-Min Chen, Sean Speese, Steve DelSignore, Tobias Stork, Mugdha Deshpande, Crystal Yu and all members of the Rodal lab for helpful discussions and technical assistance. This work was supported by NIH/NINDS (DP2 NS082I27, A.A.R.).

## Supplemental material

**Supplemental Figure 1 to Figure 2. Components of the pT-STEP targeting vector and outline of the tissue-specific tagging process of endogenous proteins in Drosophila.**
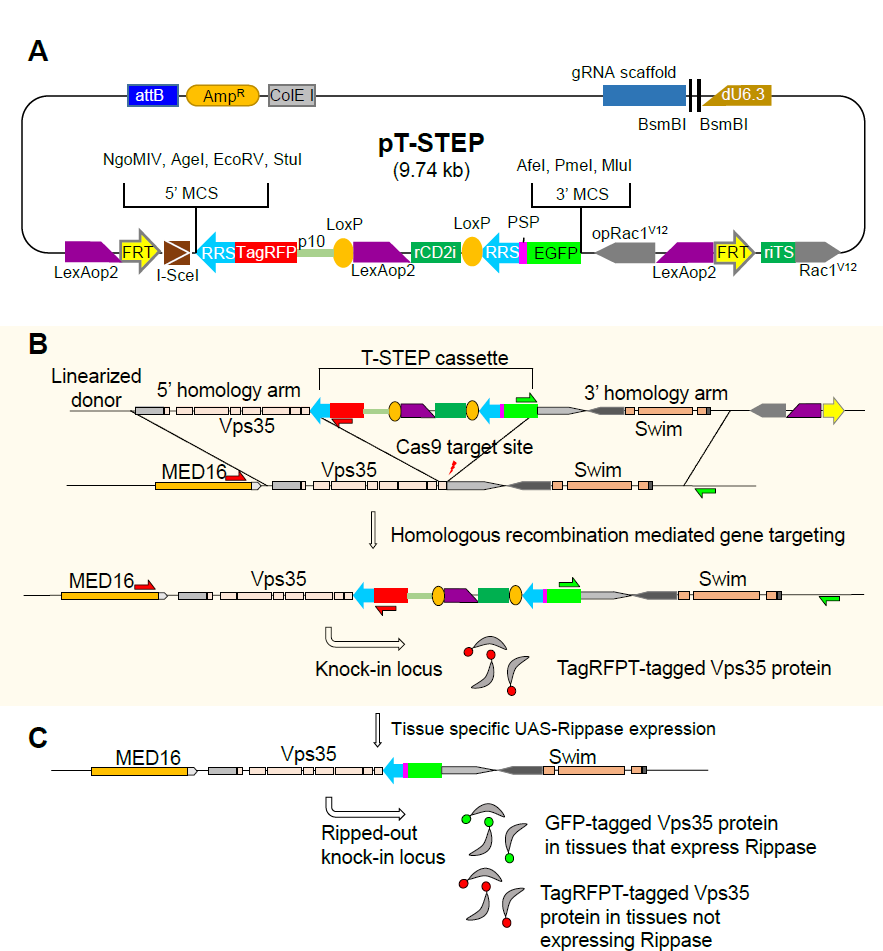
**(A)** pT-STEP vector, based on pTL2 (Chen et al. 2015), uses the same miRNA suppressor based lethality selection cassette as Golic+. BsmBI sites allow the efficient insertion of gRNA oligonucleotides, the 5’ and 3’ multiple cloning sites (MCS) are used for cloning the 5’ (in frame with TagRFPT) and 3’ homology arms, respectively. RRS, blue arrow, is the recognition sequence target of Rippase, PSP is a short recognition sequence for PreScission Protease, dU6.3 is the promoter of a small nuclear RNA U6 at 96Ac (CR31539), rCD2i is the miRNA against rat CD2, and riTS the target site for rCD2i miRNA. *p10* indicates the 3’ regulatory elements from *Autographi alifornia* nucleopolyhedrovirus. **(B)** A double stranded DNA break is induced near the C-terminus of targeted genes by Cas9 expressed in cytoblasts, as illustrated for Vps35 (gene architecture not drawn to scale). The same cytoblasts also express Flippase and I-SceI (not shown), enzymes that respectively excise and linearize the donor, which then serves as a template to repair the dsDNA break. Successful targeting events lead to TagRFPT-tagged proteins in all tissues where they are normally expressed, while the translated 5’ RRS recognition sequence serves as a small peptide linker between the tagged protein and TagRFPT. MED16 and Swim are neighboring genes. **(C)** In knock-in fly tissues that express Rippase, the DNA between the two RRS recognition sequences is excised and degraded, and a single RRS sequence remains (designed to be in frame with GFP), effectively replacing TagRFPT with a GFP tag in a tissue-specific manner.

**Supplemental Figure 2. Molecular characterization and Western blotting of T-STEP knock-in flies.**
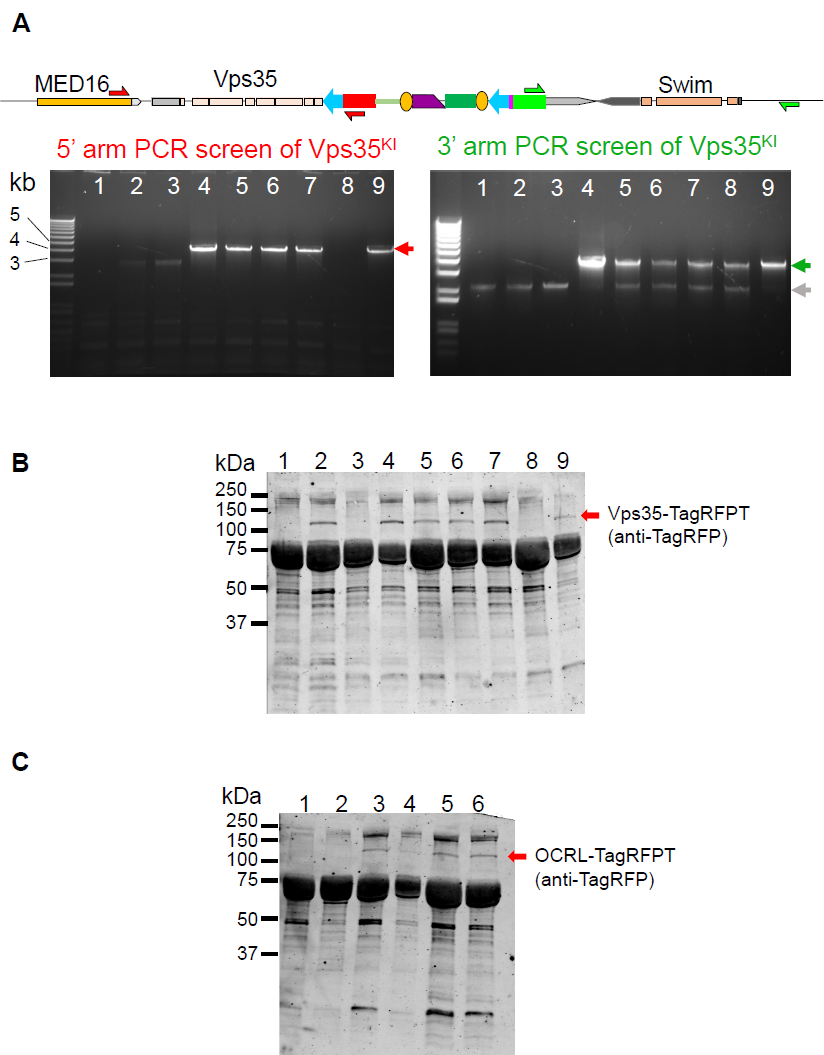
**(A)** The 5’- and 3’-most primers bind outside the region of the homology arm contained in the donor construct (also see Supplemental Figure 1 to Figure 2) and therefore diagnostic PCR yields appropriately-sized products only in cases of correctly targeted events (red and green arrows, gray arrow points to a non-specific PCR band). MED16 and Swim are neighboring genes. Genotypes are as follows: lane 1: bam898-Cas9-2A-FLP-2A-I-SceI in su(Hw)attP8/FM6c; Pin/CyO; LexAop2-5xriTS-Rac 1^V12^; lane 2: donor line Vps35-790.25.F1 in VK00027; lane 3: false positive Vps35^KI2^ where the repressor does not reside on the 2^nd^ chromosome; lane 4: Vps35 ^KI4^; lane 5: Vps35 ^KI5^; lane 6: Vps35 ^KI6^; lane 7: Vps35 KI^9^; lane 8: Vps35 KI^13^; lane 9: Vps35 ^KI16^. The 5’ PCR of Vps35 ^KI13^ line (lane 8), indicates a defective homology based repair as the TagRFPT signal on the corresponding Western blot is also absent, while the 3’ arm appears to have been incorporated during the targeting event. **(B)** Western blots with anti-TagRFP antibody of Vps35^KI^ larval hemolymph from 3.8 larvae, predicted molecular weight of Vps35-TagRFPT is 121.8 kDa. Numbers correspond to the genotypes as listed in panel A. Lane 2, the homozygous donor line residing on the 3^rd^ chromosome (in VK00027) expresses the full-length TagRFPT-tagged protein, suggesting that the regulatory elements comprised in the 3.4 kb long 5’ homology arm are sufficient to drive Vps35 expression. Note the absence of 5’ PCR product for this genotype. **(C)** Western blot with anti-TagRFPT antibody of OCRL^KI^ larval hemolymph from 3.8 larvae. The predicted molecular weight is 126.3 kDa. Lane 1: LexAop2-5xriTS-Rac1^V12^ in attP40/CyO; bam898-Cas9-2A-FLP-2A-I-SceI in attP2/TM3, Sb; lane 2: donor line OCRL-793.R39.1 in attP40; lane 3: OCRL^KI1^; lane 4: OCRL^KI3^; lane 5: OCRL^KI22^; lane 6: OCRL^KI59^.

**Supplemental Figure 3. Subcellular localization of Vps35-TagRFPT and OCRL-TagRFPT in homozygous knock-ins.**
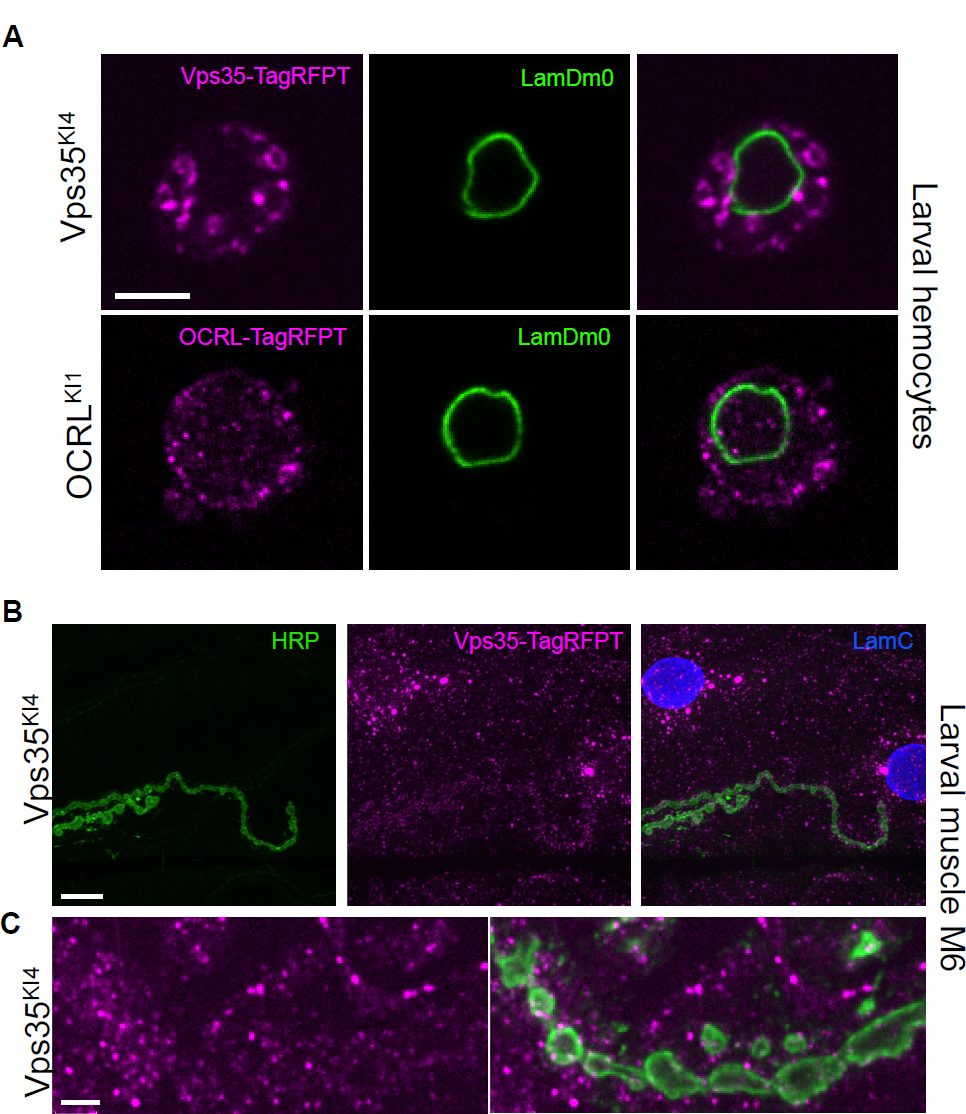
**(A)** Larval hemocytes reveal the subcellular distribution of Vps35 and OCRL, LamDm0 demarcates the nucleus. OCRL is present in smaller punctate structures, including in the nucleus, while Vps35 is frequently associated with larger, rounder endocytic compartments and not observed in the nucleus. Single confocal slices are shown, scale bar is 5 μm. **(B)** Vps35-TagRFPT localization in larval muscle 6, segment A3. LamC labels muscle nuclei. Vps35 is found throughout the muscle in small punctate structures, with more prominent accumulations near the nucleus. Maximum intensity projection of a confocal Z-stack is shown. Scale bar 10 μm. **(C)** Higher magnification view of a neuromuscular junction from another muscle 6 showing in greater detail the punctate distribution of endogenous Vps35-TagRFPT. Single confocal slice is shown, scale bar 3 μm.

**Supplemental Figure 4. Details of the available T-STEP vectors.**
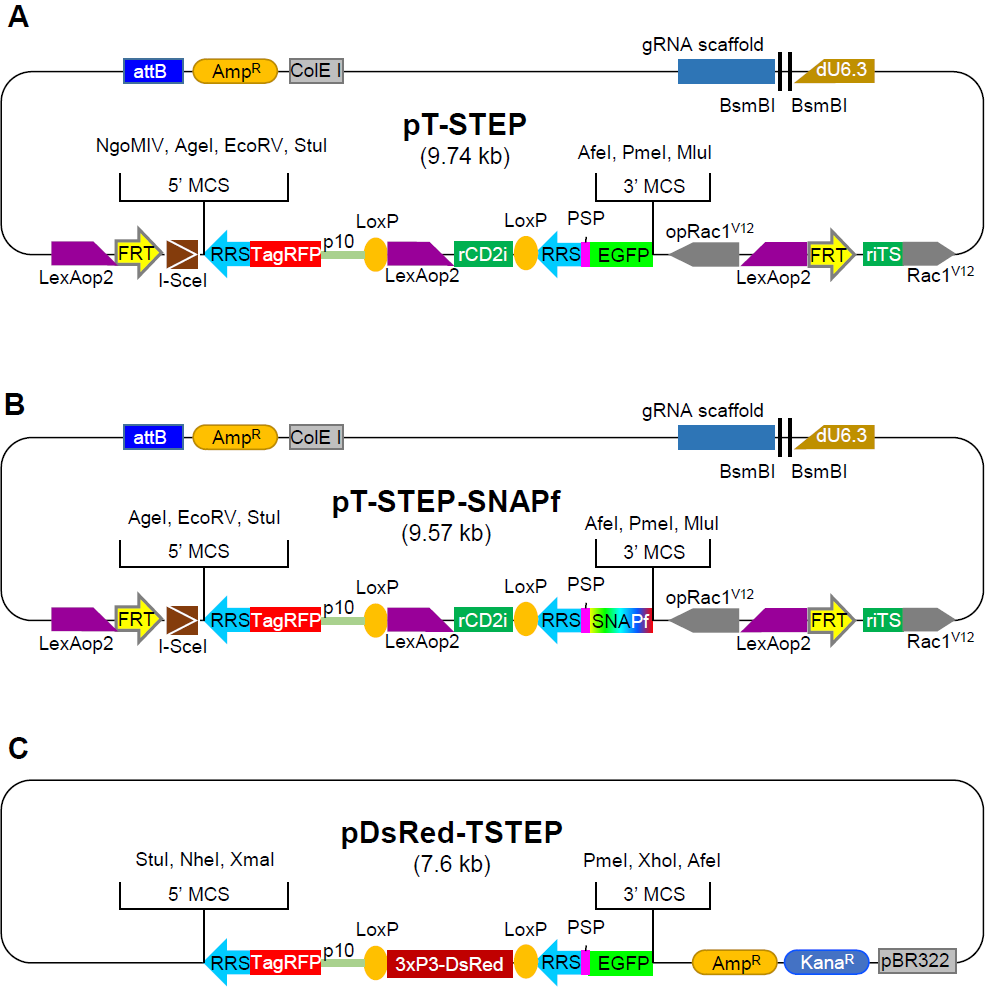
**(A)** The pT-STEP vector described and used in this manuscript for a fully transgenic T-STEP knockin approach. **(B)** pT-STEP-SNAPf differs from pT-STEP only in the presence of SNAPf instead of GFP as the swappable tag. **(C)** pDsRed-TSTEP is a targeting vector suitable for embryo injection methods, and carries DsRed to facilitate visual screening of the targeted events. After gene targeting, the tissue-specific tagging steps remain the same as described in this manuscript.

**Supplemental Figure 5. Practical details of the pT-STEP gene targeting procedure.**
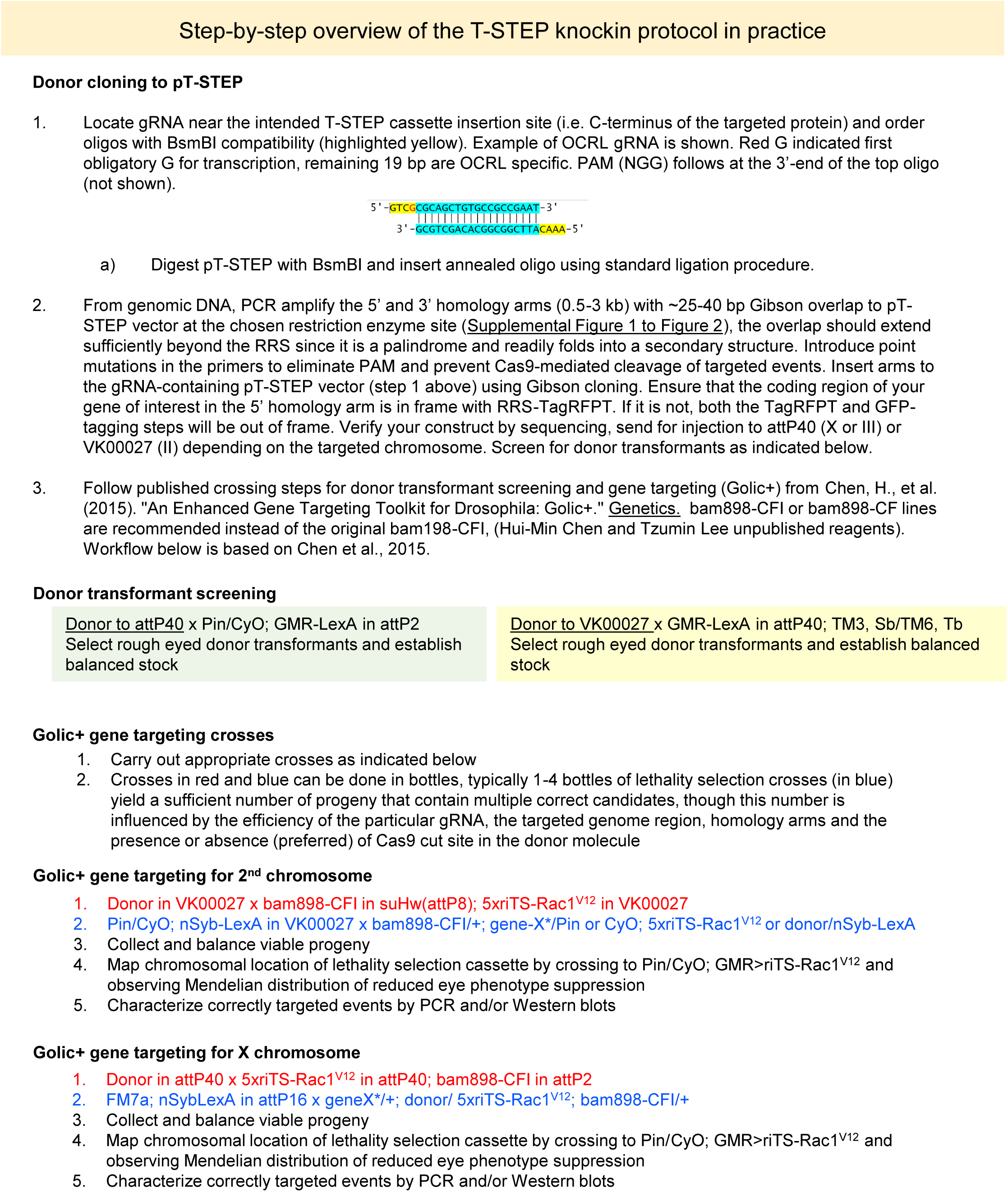
Key practical considerations include maintaining the coding frame between targeted gene and TagRFPT during cloning of the 5’ homology arm, and eliminating Cas9 cleavage of the donor construct by silencing the PAM or key residues of the Cas9 target sequence within the donor.

**Supplemental Table 1. Gene targeting efficiencies for Vps35 and OCRL.** Lines mapped refers to the number of viable progeny from the lethality selection cross that were analyzed by chromosome mapping (not all viable progeny were mapped). Targeted events indicate the number of flies where the lethality suppressor mapped to the targeted chromosome (i.e. 2^nd^ for Vps35 and X for OCRL). False positive events include escapers and local integrations, and were not characterized in detail. For Vps35D^628^N knockins, the numbers reflect targeting to the correct chromosome, and not the presence of D628N mutation in the targeted event. Experimental estimates in *Drosophila* suggest that donor utilization drops to approximately 50% within 500 bp on either side of the dsDNA cut site (Carroll and Beumer 2014), and decreases further with increasing distance. Indeed, only one of the six analyzed gene targeting candidates from the Vps35D^628^N donor construct carried the D628N mutation, which is 650 bp from the 3’ end of the 5’ homology arm. Our OCRL donor construct did not carry Cas9 resistance, unlike the Vps35 constructs, and small indels were present in several of the correctly targeted OCRL candidates, which were never observed in Vps35 targeted candidates (data not shown), arguing for the routine elimination of Cas9 target site in the donor. See also Supplemental Figure 5.

| Gene targeted | Lines mapped | Targeted events | False-positive events | Targeting efficiency |
| --- | --- | --- | --- | --- |
| Vps35 (wild type) | 14 | 12 | 2 | 86% |
| Vps35 (D628N mutant) | 10 | 6 | 4 | 60% |
| OCRL | 24 | 7 | 14 | 29% |

